# Molecular and anatomical roadmap of stroke pathology in immunodeficient mice

**DOI:** 10.1101/2022.07.28.501836

**Authors:** Rebecca Z Weber, Geertje Mulders, Patrick Perron, Christian Tackenberg, Ruslan Rust

## Abstract

**Background:** Stroke remains a leading cause of disability and death worldwide. It has become apparent that inflammation and immune mediators have a pre-dominant role in initial tissue damage and long-term recovery following the injury. Still, different immunosuppressed mouse models are necessary in stroke research e.g., to evaluate therapies using human cell grafts. Despite mounting evidence delineating the importance of inflammation in the stroke pathology, it is poorly described to what extent partial immune deficiency influences the overall stroke outcome.

**Methods:** Here, we assessed the stroke pathology of popular genetic immunodeficient mouse models, i.e., NOD scid gamma (NSG) and recombination activating gene 2 (Rag2^-/-^) mice as well as pharmacologically immunosuppressed mice and compared them to immune competent, wildtype (WT) C57BL/6J mice up to three weeks after injury. We performed histology, gene expression profiling, serum analysis and functional behavioural tests to identify the impact of immunosuppression on the stroke progression.

**Results:** We detected distinct changes in microglia infiltration, scar-forming and vascular repair in immune-suppressed mice three weeks after injury. Gene expression analysis of stroked tissue revealed the strongest deviation from immune competent mice was observed in NSG mice, for instance, affecting immunological and angiogenic pathways. Pharmacological immunosuppression resulted in the least variation in gene expression compared with the WT. Major differences have been further identified in the systemic inflammatory response following stroke acutely and three weeks following injury. These anatomical, genetic, and systemic changes did not affect functional deficits and recovery in a time course of three weeks. To determine whether the timing of immunosuppression after stroke is critical, we compared mice with acute and delayed pharmacological immunosuppression after stroke. Mice with a delayed immunosuppression (7d) after stroke showed increased inflammatory and scarring responses compared to animals acutely treated with tacrolimus, thus more closely resembling WT pathology. Transplantation of human cells in the brains of immunosuppressed mouse models led to prolonged cell survival in all immunosuppressed mouse models, which was most consistent in NSG and Rag2^-/-^ mice.

**Conclusions:** In sum, we detected distinct anatomical and molecular changes in the stroke pathology between the individual immunosuppressed mouse models that should be carefully considered when selecting an appropriate mouse model for stroke research.

## Introduction

Stroke is a major cause of disability and death due to the brain’s limited capacity to regenerate damaged tissue [1]. To promote recovery, cell-based therapy has been proposed as an emerging treatment and potential regenerative strategy for stroke patients with remaining neurologic deficits [2]. Recent advances in induced pluripotent stem cells (iPSC) technology facilitated the generation of human neuronal cells as a suitable and scalable cell source. To test the human-derived graft’s efficacy and safety for cell therapy, preclinical research requires the immunosuppression in mice to avoid xenograft rejection [3–7]. The use of immunosuppressed mice allows the evaluation of long-term effects of cell therapies in mice with the drawback that general immunosuppression may substantially alter the stroke pathology [8,9].

Acutely after stroke, inflammation plays a critical role in early ischemic damage that both promotes further injury resulting in cell death but conversely has also been shown to contribute to regeneration and remodelling [10]. Despite mounting evidence delineating the importance of inflammation in the stroke pathology, it is poorly described to what extent partial immune deficiency influences the overall stroke outcome. Several immunosuppressed mouse models have been used in preclinical stroke research, most prominent: (a) pharmacological immunosuppression with calcineurin inhibitory drugs (e.g., tacrolimus) that block the development and proliferation of T cells; (b) genetically deficient mice that lack recombination activating gene 2 protein (Rag2^-/-^) required for the generation of mature B and T lymphocytes; and (c) NOD scid gamma (NSG) mice that lack mature T and B lymphocytes, natural killer (NK) cells and have deficiencies in multiple cytokine signalling and innate immunity [11,12]. However, it is unclear to what extent the immunosuppression in these mouse models alter the natural pathology of stroke and may affect the outcome of therapies that require immunosuppression such as human cell-based therapies.

Here, we hypothesize that different aspects of stroke pathology may be altered depending on the mouse model of immunosuppression. Therefore, we compare the stroke pathology of most popular genetic mouse models using Rag2^-/-^ mice and NSG mice, as well as tacrolimus immunosuppressed mice, to immune competent, WT control. We identify changes in tissue responses and gene expression after stroke affecting especially inflammatory, scar-forming and angiogenic remodelling processes in all immunosuppressed mice compared to immune competent WT controls three weeks after injury. These changes were mitigated if the tacrolimus treatment was deferred beyond the acute stroke phase. The anatomical, genetic, and systemic changes did not affect functional deficits and recovery in a time course of three weeks. Local intraparenchymal cell transplantation led to prolonged graft survival in all immunodeficient mouse models compared to WT, which, however, was more profound in the genetically immunodeficient Rag2^-/-^ and NSG animals. These changes in anatomy, physiology, and gene expression after stroke across the tested immunodeficient mouse models are relevant to understand the role of immunosuppression in the stroke pathology and for evaluating the preclinical efficacy of cell-based therapies.

## Results

### Immunodeficient mice exhibit altered inflammation, scarring and vascular remodelling in the peri-infarct brain tissue

To identify stroke-related changes in immunodeficient mouse models, we induced a photothrombotic stroke in the right sensorimotor cortex in (1) C57BL/6J wildtype (WT) mice (2) C57BL/6J WT mice that were continuously immunosuppressed with tacrolimus (WT-Tacr); as well as genetically immunodeficient (3) Rag2^-/-^ and (4) NSG mice (**Fig. 1A**). A successful stroke was confirmed in all groups by ≈ 70% reduction of cerebral blood flow in the lesioned right hemisphere 24h after stroke induction (WT = −72%; WT-Tacr = −76%; Rag2^-/-^ = −69% NSG = −74%, all p > 0.5, **Fig. 1B, C**). Twenty-one days post injury (dpi), brain tissue was collected and revealed comparable stroke volumes and a near complete overlap (> 99%) of affected brain areas in mice from all groups (WT = 2.8 ± 0.7 mm^3^; WT-Tacr = 2.7 ± 0.7 mm^3^; Rag2^-/-^ = 2.6 ± 0.4 mm^3^; NSG = 2.4 ± 0.3 mm^3^, all p = 1, **Fig. 1D, E, Suppl. Fig. 1**). To identify stroke-related tissue responses such as inflammation, scarring and neuronal remodelling, we quantified the fluorescence intensities in the contralesional cortex as well as 300-μm cortical peri-infarct regions adjacent to the ipsilesional stroke core three weeks after injury. As expected, inflammatory as well as scarring signals were elevated in the ipsilesional compared to the contralesional cortex (**Fig. 1F-H**). In the peri-infarct areas, microglial/macrophage infiltration (Iba1^+^) and glial scarring (GFAP^+^) signals were reduced in all immunosuppressed mouse models compared to WT controls (Iba1^+^: WT-Tacr: −84%, Rag2^-/-^: −82%, NSG: −74%, all p < 0.001; GFAP^+^: WT-Tacr: −60%, p = 0.002; Rag2^-/-^: −32%, p = 0.015, NSG: −55%; p = 0.002, **Fig. 1I, J**). The quantity of mature neurons indicated by NeuN^+^ expression was reduced in the lesioned regions in all groups by 50-60% but no changes have been observed in NeuN^+^ expression between the groups (**Fig. 1K**). The expression of neurofilament light and heavy chain, major proteins of the neuronal cytoskeleton, was also altered after stroke (**Fig. 1L**). Interestingly, we observed a significantly lower loss of Neurofilament H in WT-Tacr mice (WT-Tacr: +66%, p = 0.02) but no differences in the other immunosuppressed Rag2^-/-^ and NSG mice (Rag2^-/-^: +39%, p = 0.37; NSG =+10%, p = 0.95) compared to WT control (**Fig. 1M**). No changes have also been observed between the groups for Neurofilament L expression in the injured regions (WT-Tacr: +33%, Rag2^-/-^: +23%, NSG: +2%, all p > 0.5, **Fig. 1M**).

**Fig. 1:**
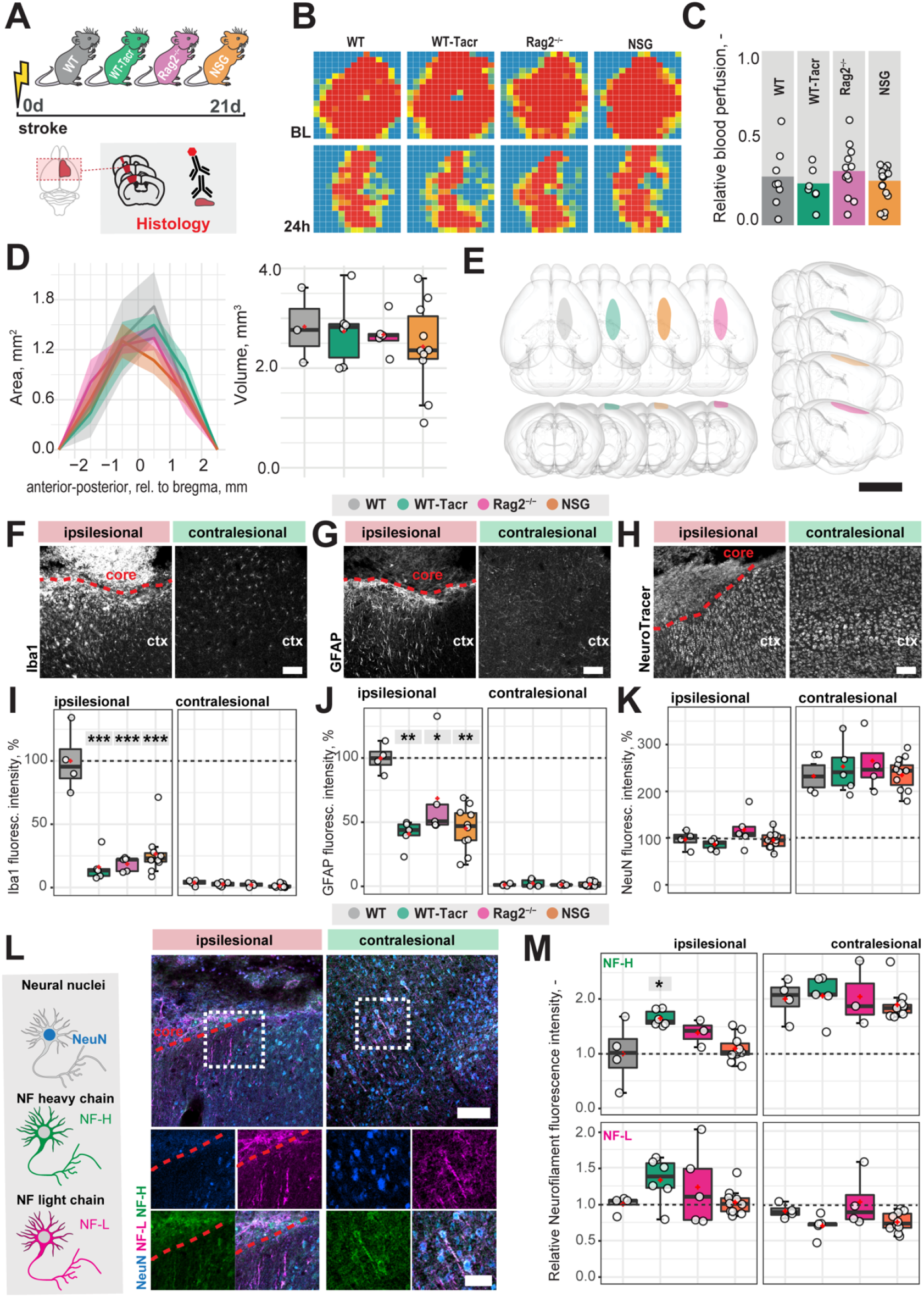
Anatomical changes in the stroke pathology of immunosuppressed mice. (**A**) Schematic representation of experimental set-up and groups of mice: C57BL/6J wildtype (WT), tacrolimus immunosuppressed wildtype (WT-Tacr), recombination activating gene 2 deficient mice (Rag2^-/-^) and NOD scid gamma mice (NSG). (B) Representative images of relative blood perfusion at baseline and 24 h after stroke induction (**C**) Quantification of cerebral blood perfusion of the injured hemisphere at 24 h following injury relative to baseline blood perfusion (**D**) Quantification of stroke area and stroke volume at 21 dpi. (**E**) 3D reconstruction of stroke location within a brain template. Scale bar: 1 cm. (**F**) Representative histological overview of activated microglia (Iba1^+^) (**G**) reactive astrocytes (GFAP^+^) and (**H**) neural nuclei (NeuroTracer) at the ipsi- and contralesional hemisphere of WT mouse. Scale bar: 100 μm. (**I**) Quantification of relative (**J**) Iba1^+^, (J) GFAP^+^ and (**K**) NeuN signal in the contra- and ipsilesional cortex 21 days following injury. (**L**) Representative histological overview of neural nuclei (NeuN^+^, blue) and neurofilament light (NF-L, magenta) and heavy chain (NF-H, green) of WT miouse. Scale bar: 50 μm. (**M**) Quantification of relative neurofilament light and heavy chain signal in the ischemic and contralesional cortex. Data are shown as mean distributions where the red dot represents the mean. Boxplots indicate the 25% to 75% quartiles of the data. For boxplots: each dot in the plots represents one animal. Line graphs are plotted as mean ± sem. Significance of mean differences between the groups (baseline hemisphere, contralesional hemisphere, and ipsilesional hemisphere) was assessed using Tukey’s HSD. Asterisks indicate significance: * P < 0.05, ** P < 0.01, *** P < 0.001. ctx: cortex, d: days, BL: baseline, dpi: days post injury, NF: neurofilament.

Next, we asked if vascular remodelling after stroke may be altered in immunosuppressed mice, because inflammatory responses are known to affect angiogenesis [13]. We quantified the vascular density and maturation as well as identified newly formed vessels in the contralesional and ipsilesional cortex three weeks after injury. The overall vascular density, number of branches and total length of the vascular network was increased in the peri-infarct areas in all immunodeficient groups compared to the WT control (**Fig. 2A-C**). Most prominent, Rag2^-/-^ mice had an increase of +70% (p < 0.001) in vascular network density, +215% (p < 0.001) in the number of branches and +79% (p < 0.001) in the vascular length compared to WT mice. Importantly, no vascular changes were observed in the contralesional brain tissue (all p >0.05, **Fig. 2D**). To confirm that the blood vessels in the ischemic border zone were newly formed, the nucleotide analogue 5-ethynyl-2’-deoxyuridine (EdU) was systemically applied at the peak of poststroke angiogenesis at day 7 (**Fig. 2E-G**). NSG mice had an increase in EdU^+^ blood vessels per mm^2^ in the ischemic border zone compared to WT controls (NSG: 109.4 ± 32.4 mm^-2^; WT: 39.2 ± 46.9 mm^-2^; p = 0.03, **Fig. 2G**). As newly formed blood vessels after stroke may be associated with immature vessels that lack stabilizing pericytes (CD13^+^) we investigated whether CD31 and CD13 markers were colocalized (**Fig. 2H, I**). Pericyte coverage in the ipsilesional site was ≈ 70% lower compared to the contralesional site in all groups but was comparable between the groups in the periinfarct areas as indicated by the ratio of pericyte covered vasculature (CD 13^+^CD31^+^) to total vasculature (CD31^+^) (WT: 0.36 ± 0.15, WT-Tacr: 0.15 ± 0.14, Rag2^-/-^: 0.23 ± 0.12, NSG = 0.23 ± 0.1; all p > 0.4, **Fig. 2J**).

**Fig. 2:**
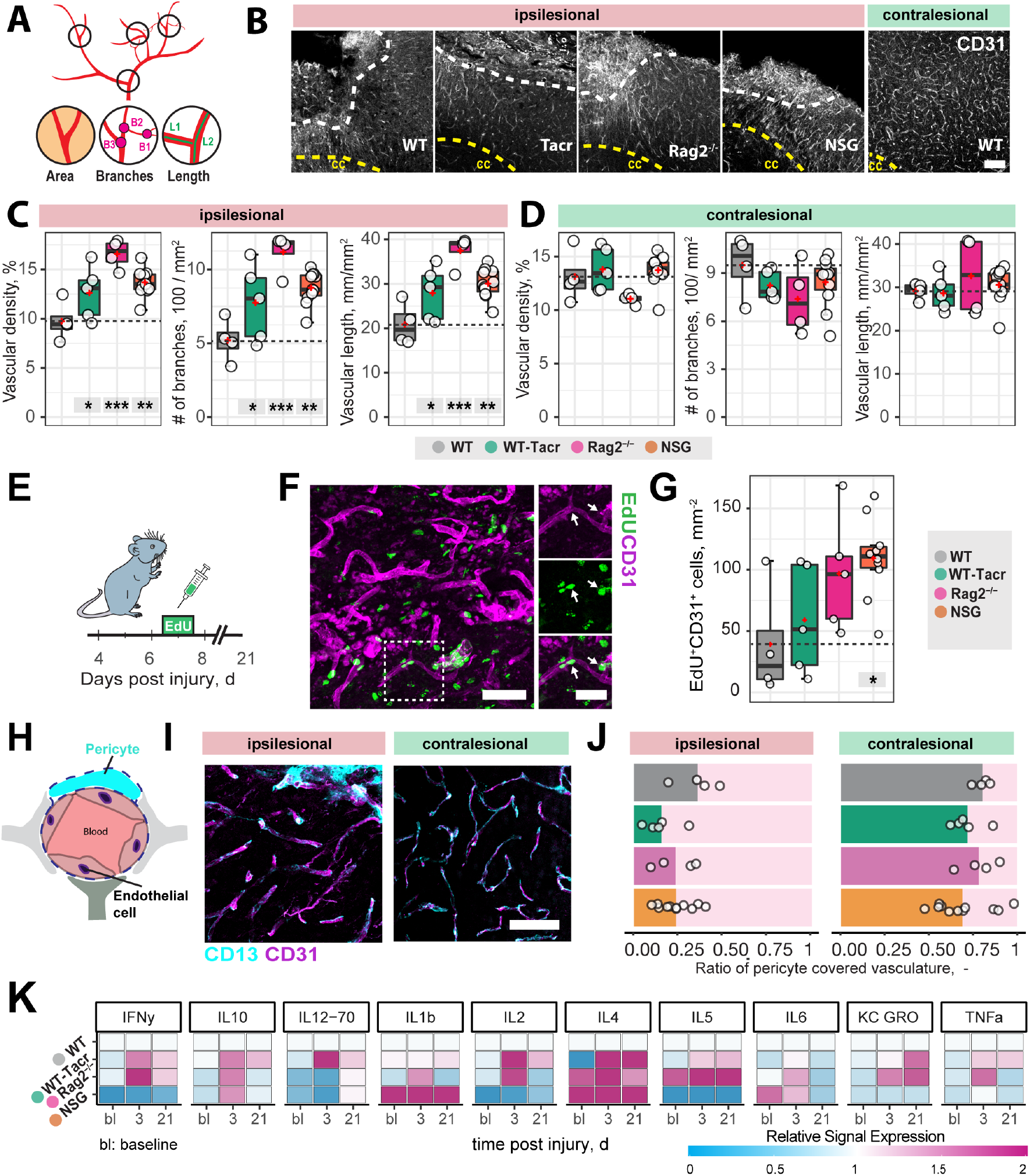
Vascular repair and remodelling of immunosuppressed mice after stroke. (**A**) Schematic representation of vascular parameters (**B**) Representative images of CD31^+^ vasculature in the ipsi- and contralesional cortex in WT mouse. Scale bar: 100 μm. (**C**) Quantification of vascular density, number of vascular branches and vascular length of ipsi- and (**D**) contralesional cortex 21 dpi. (**E**) Schematic representation of EdU injection to label proliferating cells 7 dpi. (**F**) Representative image of EdU incorporating CD31 ^+^ blood vessels in the ipsilesional site at 21 dpi. Arrow shows EdU^+^ vasculature (**G**) Quantification of EDU^+^CD31^+^ cells per mm^2^ in the ipsilesional site. (**H**) Schematic representation of blood brain barrier components. (I) Representative image of CD31^+^ cells covered by CD13^+^ pericytes in the ipsi-and contralesional site of WT mice at 21 dpi. Scale bar: 25 μm. (**J**) Quantitative ratio of vasculature covered by pericytes in ipsi- and contralesional site. (**K**) Blood plasma cytokine analysis of pro-inflammatory factors at baseline, 3 and 21 dpi. Data are shown as mean distributions where the red dot represents the mean. Boxplots indicate the 25% to 75% quartiles of the data. For boxplots: each dot in the plots represents one animal. Significance of mean differences between the groups (baseline hemisphere, contralesional hemisphere, and ipsilesional hemisphere) was assessed using Tukey’s HSD. Asterisks indicate significance: * P < 0.05, ** P < 0.01, *** P < 0.001. ctx: cortex, bl: baseline, d: days.

To test if also systemic inflammatory responses were altered following stroke, we measured serum cytokine levels at baseline, 3 and 21 days after stroke using MSD multiplex assays (**Fig. 2K**). We identified that all immunosuppressed mice showed a changed cytokine signature compared to WT mice. The strongest deviation from WT was present in cytokine levels of NSG mice, particularly in the reduced levels of IFNy, IL-2, IL-5 and increased Il-1b, Il-4 (**Fig. 2K**).

### Motor performance and gait alterations after stroke are comparable between immunodeficient and wildtype mice

Large damage of the sensorimotor cortex by photothrombotic stroke causes long-term impairment in fore and hindlimb function and locomotion [14,15]. To test whether the anatomical and systemic changes following immunosuppression may affect functional recovery, motor performance and gait analysis was performed at baseline and at repeated intervals after stroke for 21 days (**Fig. 3A**).

**Fig. 3:**
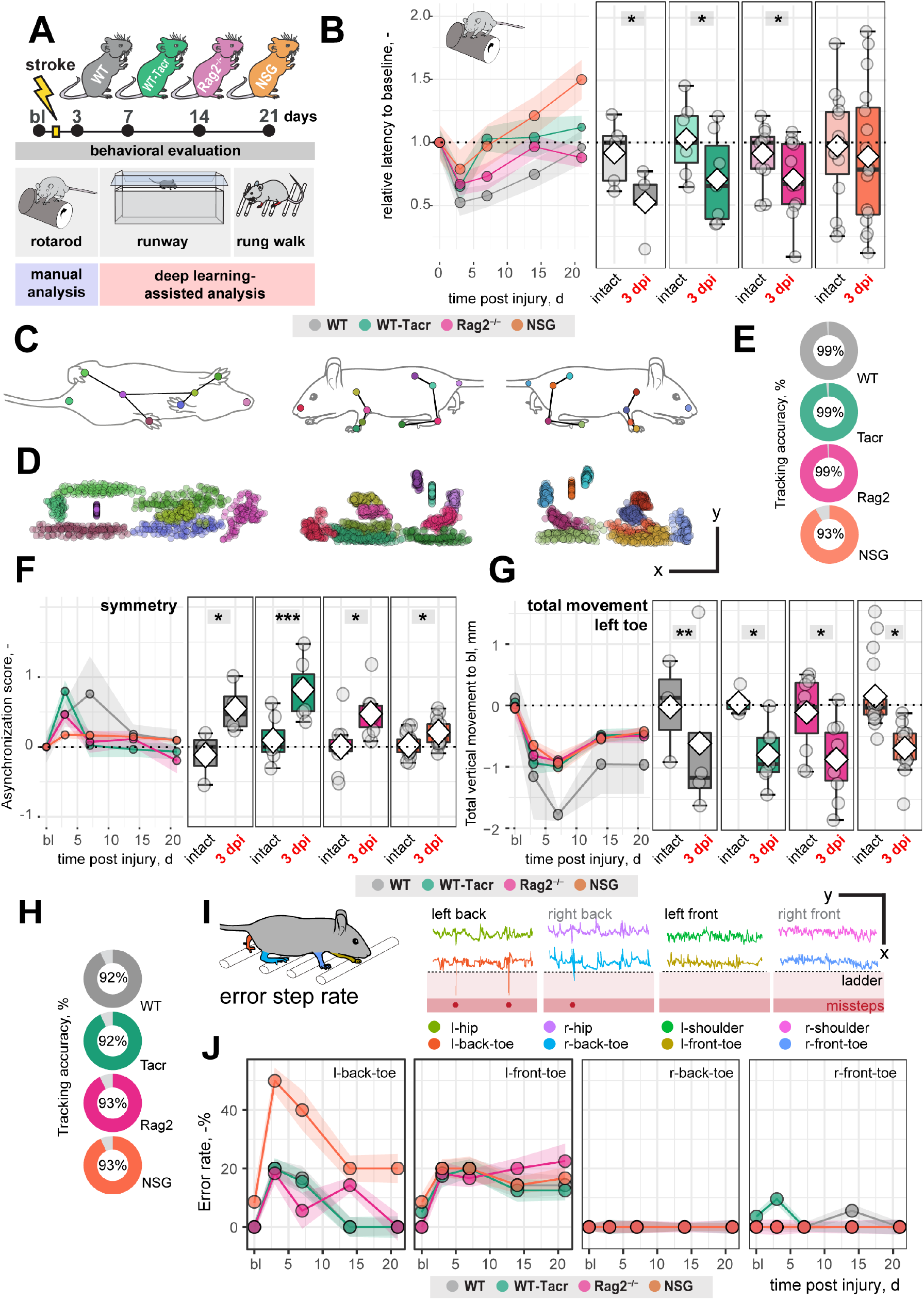
Gait and motor performance of immunosuppressed mice after stroke. (**A**) Schematic time course of experimental interventions (**B**) Rotarod test showing the relative latency at baseline and 3, 7, 14, 21 dpi. (**C, D**) Walking profile normalized to the hip coordinates of randomly selected non-injured mice. Each dot represents an anatomical landmark. (**E**) Likelihood of a confident labelling for individual body parts in the runway (**F**) Ratio of asynchronization at baseline, 3,7,14,21 dpi. (0 = ideal synchronization). (**G**) Absolute vertical movement of left (contralesional) toe at baseline and 3,7,14,21 dpi. (**H**) Likelihood of a confident labelling for individual body parts in the ladder rung test (**I**) Representation of an individual ladder rung walk of WT mice at 3 dpi, selected body parts are tracked including hip, back toe, shoulder, and front toe. (**J**) Time course of error rate during ladder rung test in the individual paws. Data are shown as mean distributions where the white dot represents the mean. Boxplots indicate the 25% to 75% quartiles of the data. For boxplots: each dot in the plots represents one animal. Line graphs are plotted as mean ± sem. For line graphs: the dots represent the mean of the data. Significance of mean differences between the groups was assessed using repeated ANOVA with post-hoc (emmeans) analysis. Asterisks indicate significance: * P < 0.05, ** P < 0.01, *** P < 0.001.

We determined in all groups, except of NSG mice, an acute deficit in the rotarod performance indicated by a ≈ 40% shorter time spent on the rod acutely after stroke (3dpi to bl: WT: −48%, p = 0.020, WT-Tacr: −35%, p = 0.048; Rag2^-/-^ = −43%, p = 0.037; NSG = −22%, p = 0.549) with a gradual recovery (**Fig. 3B**). NSG mice had also a striking higher intra-group variability between individual animals in the rotarod task (**Fig. 3B**).

Next, we applied a recently established deep learning algorithm to videos of mice during a voluntary run to identify specific gait changes after stroke [16,17]. We confirmed that the body parts of interest including tail base, iliac crest, hip, back ankles, back toe, shoulder, wrist, elbow, front toe, and head could be reliably detected from three perspectives (**Fig 3C, D**). The ratio of confident labels (>95% likelihood of detection) to total labels ranged from 93-99% (**Fig. 3E**). Data points that did not pass the likelihood of detection of 95% were excluded. We confirmed that previously identified key parameters [16] were altered in all groups of animals after stroke. For instance, an asymmetric stride pattern describing the non-simultaneous placement of opposed front and back toes, was observed acutely after stroke in all groups (3dpi: all p < 0.05) (**Fig. 3F**). The asymmetric stride pattern reversed to baseline with time in all groups (**Fig. 3F**). Also, we detected comparable irreversible gait changes in all groups such as a reduced range of motion in the contralesional left front paw (3dpi to bl: WT: −1.2 ± 1.4 mm, WT-Tacr: −0.9 ± 0.5 mm, Rag2^-/-^: −0.8 ± 0.6 mm, NSG: −0.7 ± 0.3 mm, all p < 0.05) that gradually recovered but did not reach baseline levels throughout the time course (21dpi to bl: WT: −1.0 ± 0.8 mm, WT-Tacr: −0.5 ± 0.3 mm, Rag2^-/-^: −0.5 ± 0.4 mm, NSG: −0.4± 0.3 mm, all p < 0.05, **Fig 3G**).

To detect fine motor deficits, we additionally performed a deep learning assisted irregular ladder rung test and quantified the ratio of stepping errors [16,18]. We achieved a high ratio of confident labels of 92-93% (**Fig. 3H**) and excluded all data points that did not pass the confidence threshold. Errors were defined if the front or back toe dropped lower than the ladder height due to a misstep or a slip (**Fig. 3I**). All groups of mice showed acutely increased error rates that were most pronounced in the contralateral left back (3dpi: WT: +20%, WT-Tacr: +20%, Rag2^-/-^: +18%, NSG: +50%, all p < 0.05) and left front toe (3dpi: WT: +20%, WT-Tacr: +10%, Rag2^-/-^: +18%, NSG: +18%, all p < 0.05) compared to baseline misstep rates (**Fig. 3J**). NSG mice tend to have a higher overall error rate, most prominent in the left back toe after injury (**Fig. 3J**).

In sum, the motor performance and gait alterations between immunosuppressed and WT mice were largely comparable. NSG mice deviated in few behavioral readouts from the functional deficits in WT mice. However, the data indicates that the observed anatomical and systemic changes did not lead to major changes in functional recovery between the groups in a time course of three weeks.

### Immunodeficient mice have distinct gene expression profile after stroke

To further dissect molecular changes in the ischemic brain, we performed RNA sequencing of ischemic brain tissue from immunodeficient and WT mice (**Fig. 4A**). Three weeks after stroke, the majority of the significantly altered genes were upregulated in all groups of mice (**Fig. 4B**). Principal component analysis and a heatmap of the most differentially expressed genes demonstrate a clear separation between naïve mice from their stroked littermates (**Fig. 4B, C**). As expected, we also observed a strong separation between non-stroked and stroked NSG mice to the other groups presumably because of their different background (NSG mice have a NOD-background; all other groups of mice had a C57Bl/J background, **Suppl. Fig. 2 A, B**). Most enriched gene sets between non-stroked NSG and WT mice affected pathways related to the adaptive immune system (**Suppl. Fig. 3**).

**Fig. 4:**
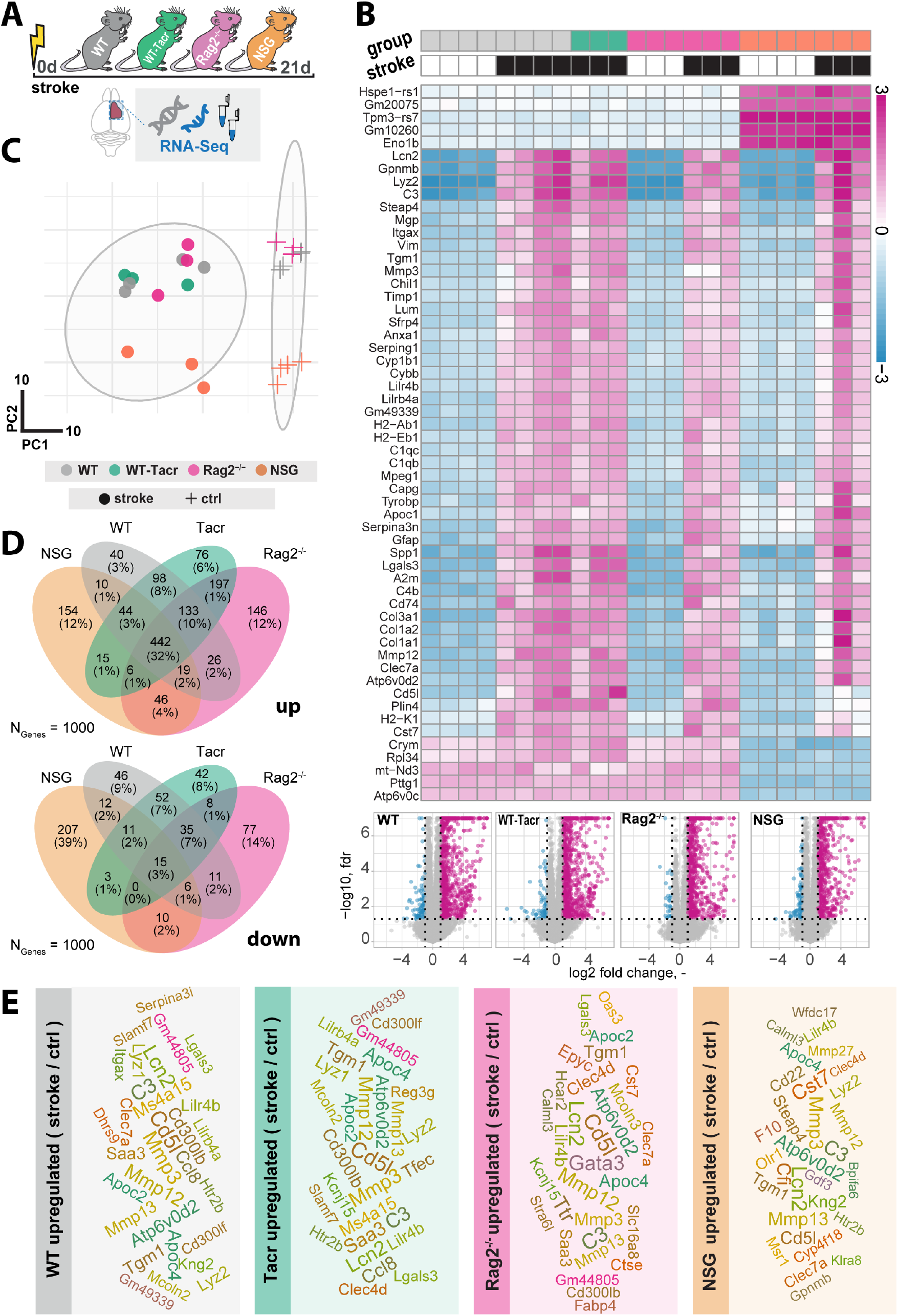
Gene expression changes in immunodeficient mouse models after stroke. (**A**) Schematic overview of experimental time course and groups of mice: C57BL/6J wildtype (WT), tacrolimus immunosuppressed wildtype (WT-Tacr), recombination activating gene 2 deficient mice (Rag2^-/-^) and NOD scid gamma mice (NSG). (**B**) Heatmap of most differentially expressed genes and volcano plot of stroked (black boxes) to non-stroked (white boxes) WT, WT-Tacr, Rag2^-/-^, NSG mice. (**C**) Principal component analysis, circles indicate stroked mice, crosses indicate non-stroked mice. (**D**) Venn diagram of common and differentially expressed top1000 genes in all groups of immunodeficient mice. (**E**) List of top 30 upregulated genes after stroke in the respective group of immunodeficient mice, font size represents strength of upregulation

After stroke, there was an overlap of 32% of the top_1000_ upregulated genes but only 3% of the top_1000_ downregulated genes in all groups of mice compared to their respective non-stroke controls (**Fig. 4.D**). WT-Tacr mice showed the closest gene expression profile to WT; and NSG mice deviated most from WT gene expression after stroke. Still, many of the top_30_ upregulated genes were present in all groups of animals such as Mmp12, CD51, Mmp3, C3, Apoc4, Tgm1, Clec7a, Lcn2 and Lilr4b. (**Fig 4E**). However, no top_30_ downregulated genes were present in all groups (**Suppl. Fig. 4**)

The majority of the enriched pathways in all groups after stroke were related to the innate and adaptive immune system e.g., leukocyte migration, positive regulation of cytokine production, and regulation of immune effector process (**Fig. 5A, B**). While 80% of the top 10 enriched pathways after stroke overlapped between WT-Tacr, Rag2^-/-^ and WT, only 40% of pathways were identical between NSG and WT (**Fig. 5A, B**).

**Fig. 5:**
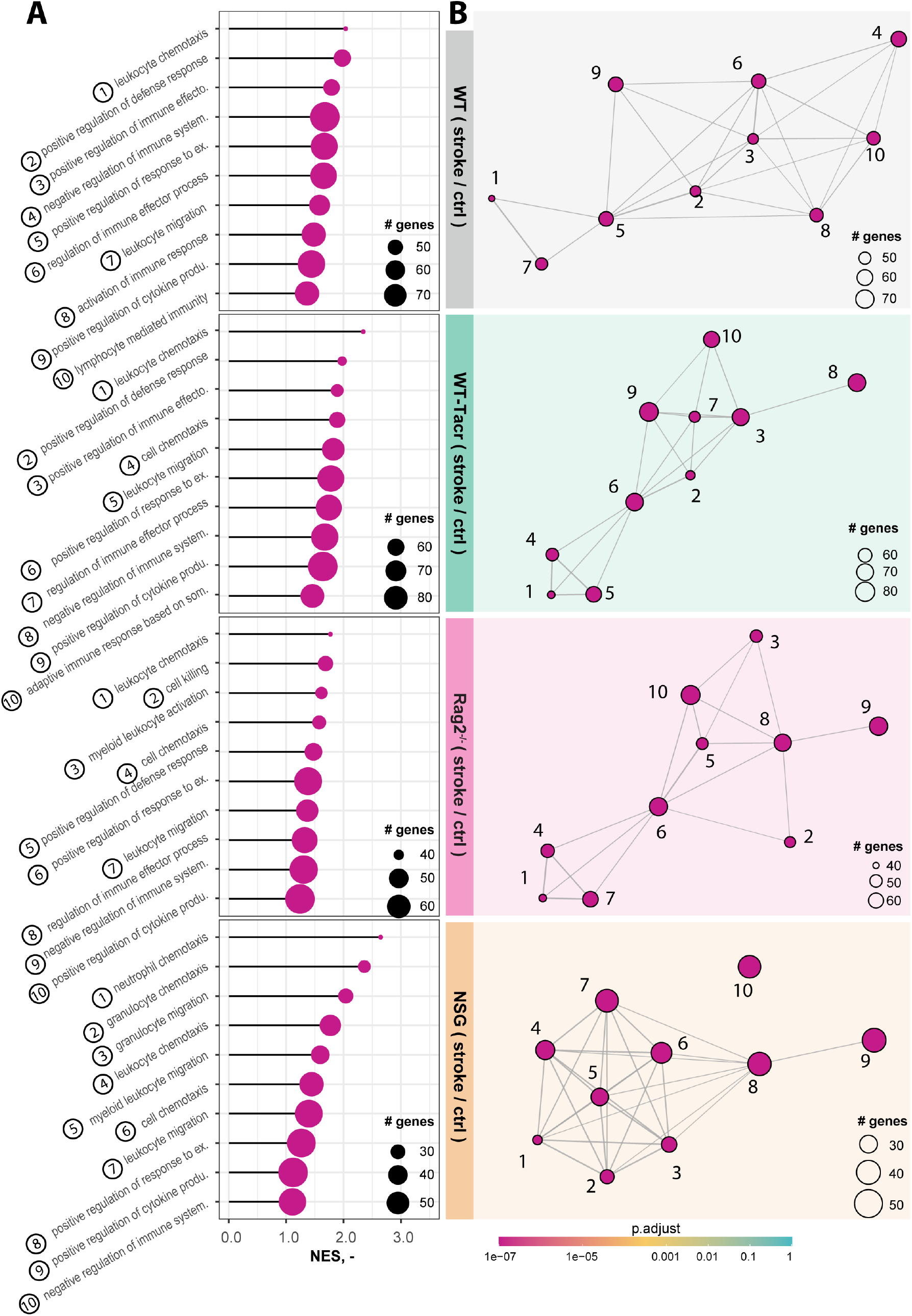
Gene set enrichment analysis in stroked mice. **(A)** Lollipop plot and (**B**) enrichment map (EMAP) of top 10 pathways enriched in stroked C57BL/6J wildtype (WT), tacrolimus immunosuppressed wildtype (WT-Tacr), recombination activating gene 2 deficient mice (Rag2^-/-^) and NOD scid gamma mice (NSG).mice compared to their non-stroked littermates. Size of dots represents the number of genes in the pathway and color of dots represents adjusted p value.

Next, we analysed gene expression differences among the immunosuppressed stroked groups that were normalized for their respective non-stroked genotype to WT stroked mice (**Fig. 6A**). The majority of differentially expressed genes in the different immunosuppressed stroked mice were unique (**Fig. 6B**). As expected, the enriched pathways were mostly associated with inflammatory responses such as, innate immune response, regulation of cytokine production and lymphocyte activation (**Suppl. Fig. 5A-C**).

**Fig. 6:**
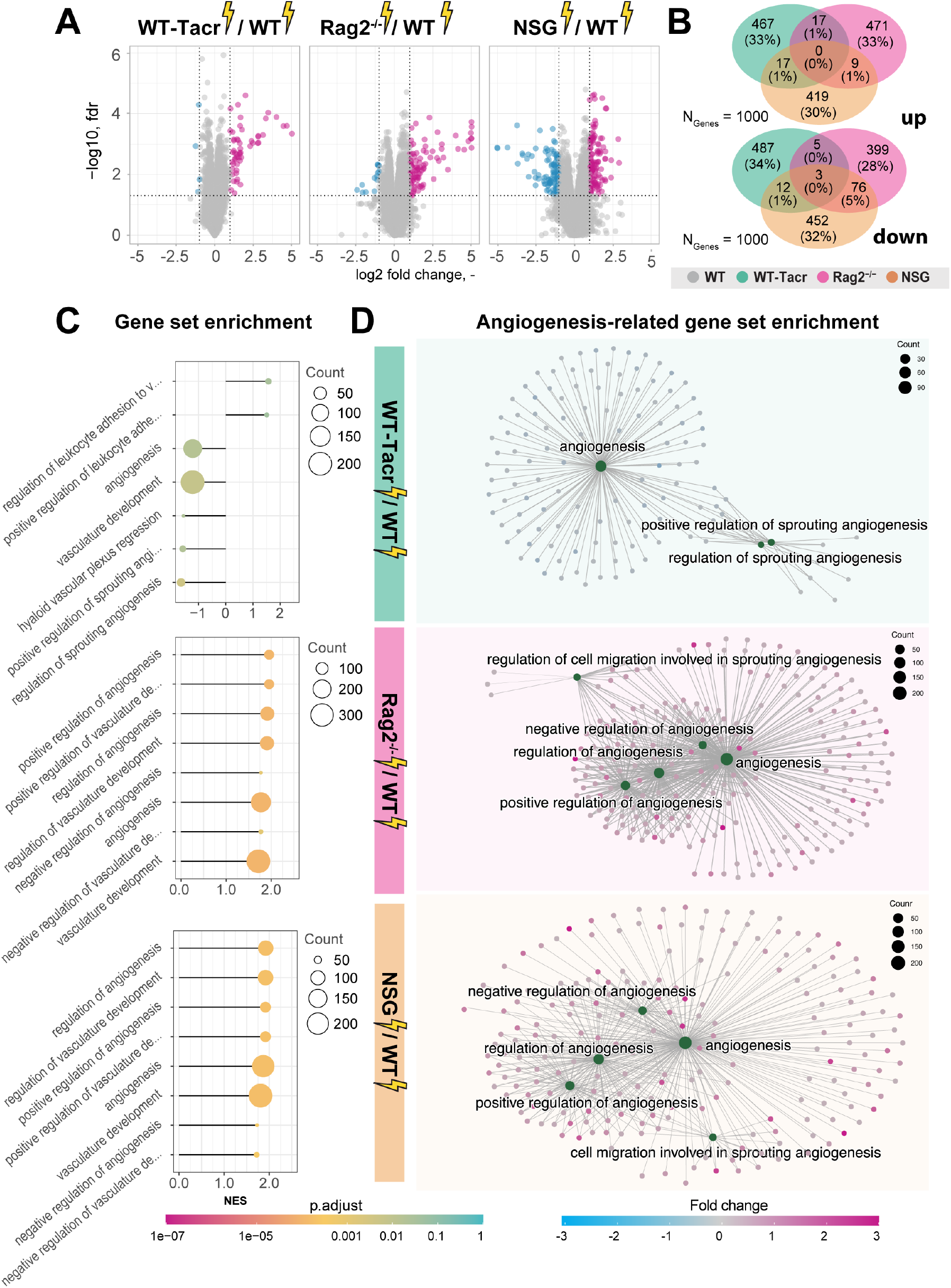
Gene expression changes and gene set enrichment between immunodeficient stroked mice and wildtype stroke pathology. (**A**) Volcano plot of differentially expressed genes identified between immunodeficient stroked mice and wildtype stroked mice. Magenta dots indicate upregulated gene expression, and blue dots indicate downregulated gene expression. (**B**) Venn diagram of common and differentially expressed top1000 genes in all groups of stroked immunodeficient mice. (**C**) Gene set enrichment shown in a lollipop plot in immunodeficient stroked mice compared to wildtype stroked mice. Size of dots represents the number of genes in the pathway and color of dots represents adjusted p value. (**D**) CNET plot describing linkages of genes and biological processes for angiogenesis-related pathways. Size of dots represents the number of genes in the pathway and color of dots represents fold changes.

Interestingly, a second cluster of pathways that was enriched in immunosuppressed stroked mice compared to WT stroked mice was linked to angiogenic responses (**Fig 6 C, D**). Surprisingly, these pathways were highly positively enriched in NSG and Rag2^-/-^ mice (**Fig. 6D**) but negatively enriched in WT-Tacr mice at 3 weeks after injury (**Fig. 6D**), indicating a reduced angiogenic response at 21 dpi in tacrolimus immunosuppressed mice.

In sum, we observed distinct gene expression and pathway enrichment changes between the immunosuppressed and WT mice three weeks after stroke. The strongest deviation was observed in NSG mice. Apart from inflammatory responses, angiogenesis related genes were most significantly altered.

### Delayed immunosuppression results in altered stroke progression that more closely resembles wildtype pathology

Transient angiogenic responses have been previously reported after stroke [19] and may explain the enhanced vascular repair mediated by tacrolimus with concomitant downregulation of angiogenesis related genes and pathways three weeks after stroke. We therefore asked if the anatomical and gene expression differences in the stroke pathology between tacrolimus treated and WT mice could be reduced by delaying the tacrolimus administration by one week. To test this hypothesis, we induced a photothrombotic stroke in (1) WT, (2) acute (0 dpi) tacrolimus immunosuppressed mice (WT-Tacr_acute_) and (3) delayed (7 dpi) tacrolimus immunosuppressed mice (WT-Tacr_delayed_, **Fig. 7A**).

**Fig. 7:**
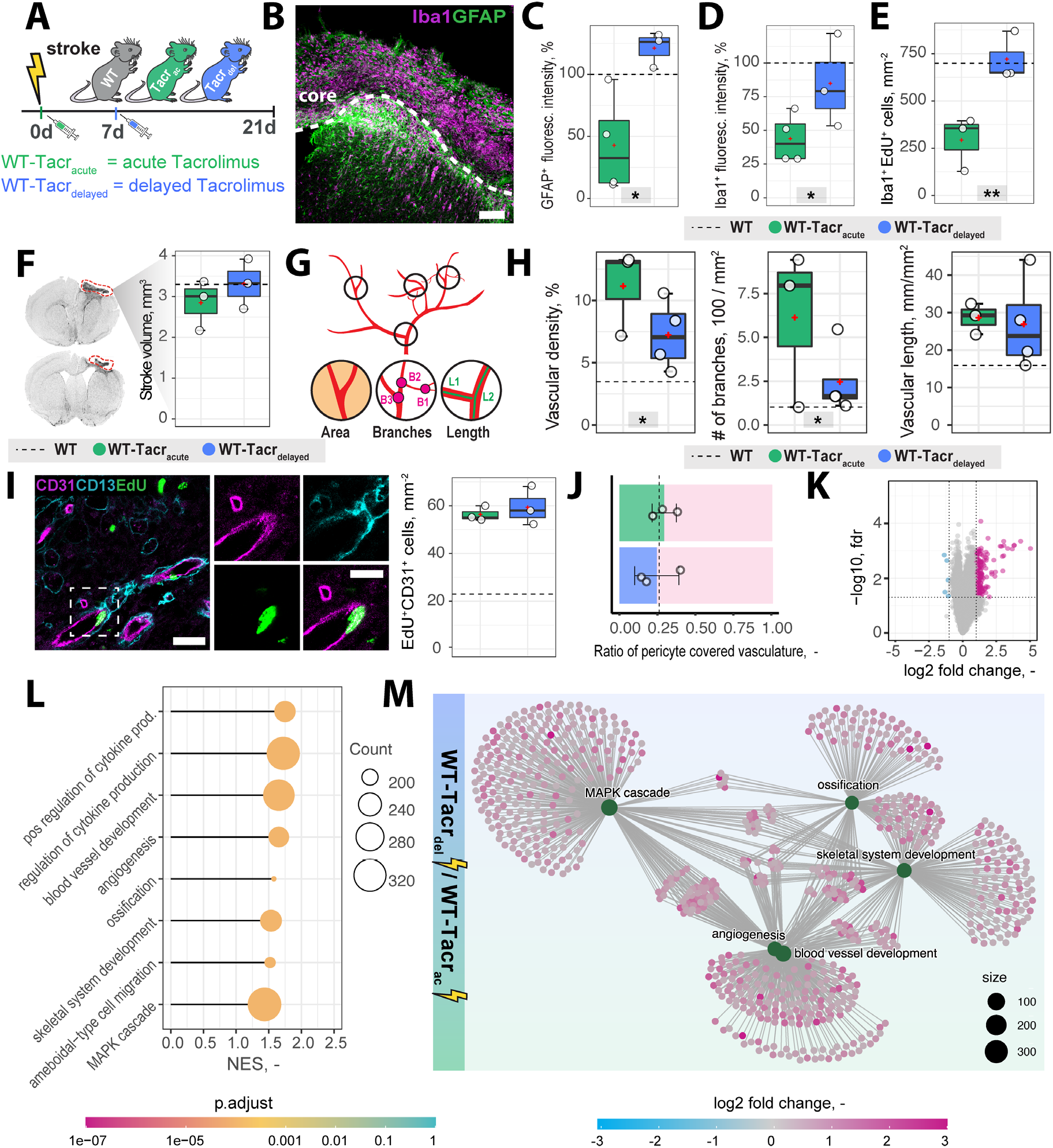
Anatomical and gene expression changes in the stroke pathology of acute and delayed pharmacological immunosuppression: (**A**) Schematic representation of experimental set-up and groups of mice: C57BL/6J wildtype (WT), acute (0 dpi) tacrolimus immunosuppressed wildtype (WT-Tacr_acute_), and delayed tacrolimus (7 dpi) immunosuppressed wildtype (WT-Tacr_delayed_) mice. (**B**) Representative image of Iba1^+^ inflammatory cells (Iba1, magenta) and reactive astrocytes (GFAP^+^, green) in the ipsilesional cortex at 21 dpi. Scale bar: 100 μm. (**C**) Quantification of GFAP^+^, and (**D**) Iba1^+^ fluorescence intensity in the peri-infarct regions at 21 dpi. (**E**) Quantification of Iba1^+^ EDU^+^ cells in per-infarct region. (**F**) Stroke volume quantification at 21 dpi. (**G**) Schematic overview of vascular parameters (H) Quantification of vascular density, number of vascular branches and vascular length in the peri-infarct region at 21 dpi. (**I**) Representative image of newly formed blood vessels (CD31^+^, magenta) covered by pericytes (CD13^+^, cyan) and incorporating a nucleotide analogue that was injected at 7 dpi (EdU^+^, green). Scale bars: Overview: 20 μm. Closeup: 10 μm. Quantification of EdU^+^CD31+ cells per mm^2^ in the peri-infarct region at 21 dpi. (**K**) Volcano plot of gene expression differences between delayed and acute immunosuppressed mice. (**L**) Lollipop chart of gene set enrichment terms for “biological process” in delayed tacrolimus immunosuppressed mice. (**M**) Visualization of top five gene set enrichments terms with respective genes. Data are shown as mean distributions where the white dot represents the mean. Boxplots indicate the 25% to 75% quartiles of the data. For boxplots: each dot in the plots represents one animal. Dotted line represents mean of WT mice.

First, we performed brain tissue analysis of inflammatory and scarring responses at 21 dpi. (**Fig. 7B**). Histological analysis in the peri-infarct region revealed an increased glial scar signal (GFAP^+^) in WT-Tacr_delayed_ (+21% to WT) compared to WT-Tacr_acute_ (−58% to WT) mice (p = 0.025, **Fig. 7C**). Inflammatory Iba1^+^ levels were also amplified in WT-Tacr_delayed_ (−25% to WT) compared to WT-Tacr_acute_ (−60% to WT) mice (p = 0.04, **Fig. 7D)**. The number of newly formed Iba1^+^ cells (Iba1^+^EdU^+^) around the stroke was increased between Tacr_delayed_ (+1% to WT) compared to WT-Tacr_acute_ (−59% to WT) mice (p = 0.001, Fig. 7E), indicating overall that post-stroke scarring and inflammation in WT-Tacr_delayed_ mice is closer to WT tissue responses.

No differences have been observed in the stroke volumes between the groups (WT-Tacr_acute_ = 2.85 ± 0.62 mm^3^, WT-Tacr_delayed_ = 3.31 ± 0.61 mm^3^, WT = 3.30 ± 0.71 mm^3^, p >0.05, **Fig. 7F**). Additionally, delayed tacrolimus administration reduced the vascular density (−35%, p = 0.048) and number of branches (−60%, p = 0.032) but did not influence the number of newly formed blood vessels compared to WT-Tacr_acute_ (CD31^+^EDU^+^: WT-Tacr_acute_ = 56.3 ± 3.2 cells/mm^2^, WT-Tacr_delayed_ = 59.3± 8.1 cells/mm^2^, p > 0.05, **Fig. 7 G-I**). The ratio of pericyte coverage of total vasculature remained unchanged in the peri-infarct region between the groups (WT-Tacr_acute_ = 0.25 ± 0.15, WT-Tacr_delayed_ = 0.29 ± 0.08, p > 0.05, **Fig. 7J**).

Interestingly, gene expression analysis of ischemic brain tissue reveals an increased gene set enrichment for cytokine production and pro-angiogenic responses in WT-Tacr_delayed_ mice (**Fig 7L, Suppl. Fig. 6**). This may indicate that the increased angiogenic responses in WT-Tacr_acute_ mice may be transient and/or that delayed immunosuppression shifts the angiogenic response towards a later timepoint after stroke.

These data indicate that a 7-day delayed pharmacological immunosuppression with tacrolimus shifts the course of stroke more towards the WT pathology.

### Grafted human NPCs survive long-term in genetically immunodeficient Rag2^-/-^ and NSG mice

Next, we evaluated long-term survival of intracerebrally transplanted grafts in (1) WT mice (2) WT mice that were continuously immunosuppressed with tacrolimus (WT-Tacr); and genetically immunodeficient (3) Rag2^-/-^ and (4) NSG mice (**Fig. 8A**). As a cell source, we used recently generated human neural progenitor cells (NPCs) derived from induced pluripotent stem cells (iPSCs) [20]. The NPCs were transduced with a dual reporter consisting of a bioluminescent firefly luciferase (rFluc) and a fluorescent eGFP. A successful transplantation in all groups of mice was confirmed by comparable levels of luciferase signal directly after transplantation (**Fig. 8B, C**). As expected, we observed the absence of the graft in WT mice 14 days after transplantation presumably due to xenogenic immune rejection. In contrast, the immunosuppressed mice showed all a prolonged graft survival. However, only in Rag2^-/-^ and NSG mice, we observed a strong bioluminescence signal during the entire time course of more than 1 month after transplantation (**Fig. 8B, C**). After 35 days, surviving graft cells were identified using human-specific antibodies in brain sections of NSG and Rag2^-/-^ mice but were not detectable in WT and WT-Tacr mice (**Fig. 8D**).

**Fig. 8:**
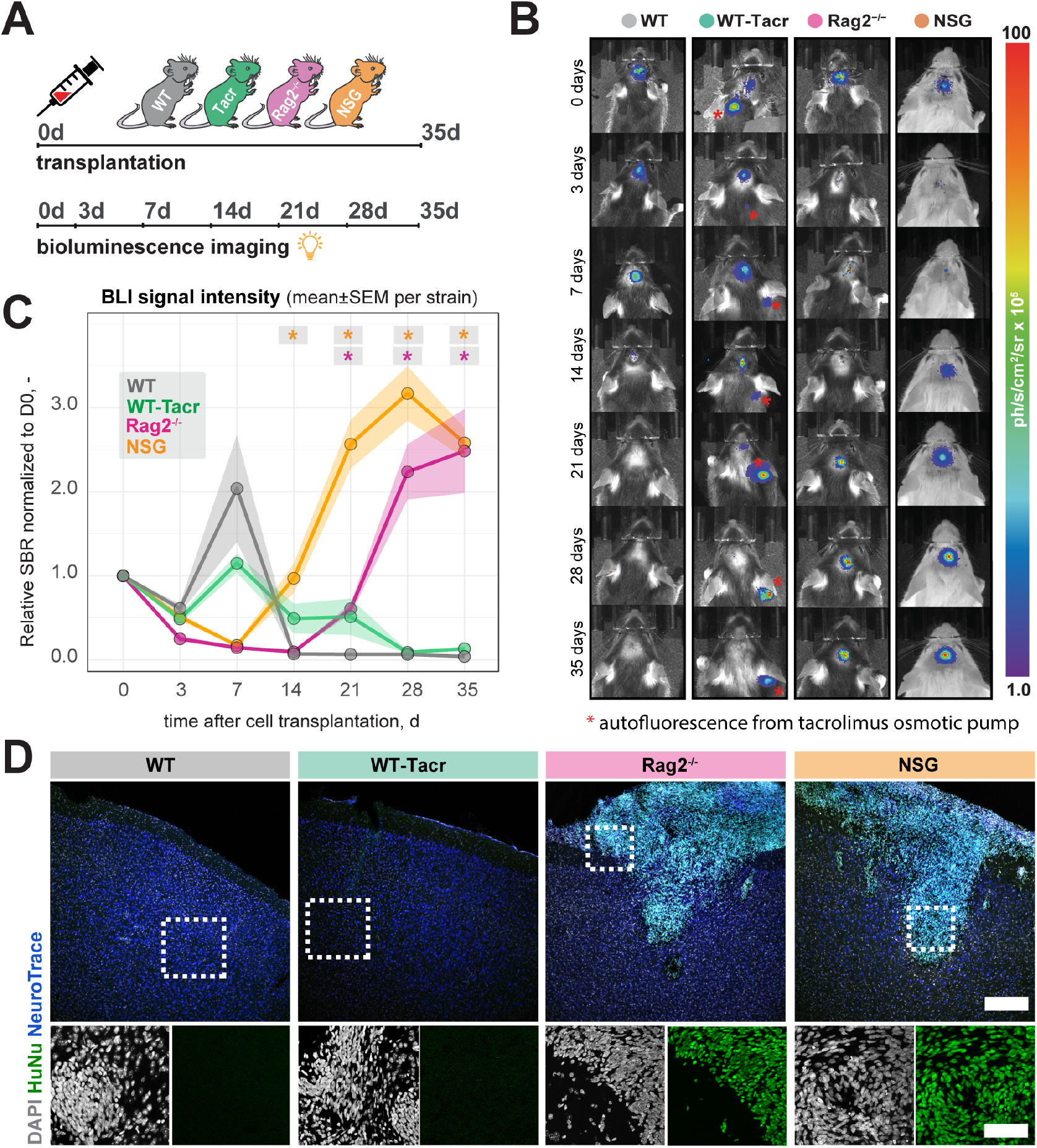
*In vivo* tracking of transplanted NPCs in immunodeficient mouse models. (**A**) Schematic overview of experimental time course (**B**) Representative bioluminescence images of luciferase-expressing NPCs transplanted in immunodeficient and WT mouse models at 0, 3, 7, 14, 21, 28, 35 days after transplantation. (**C**) Bioluminescence signal intensity over time after transplantation. (D) Representative images of transplanted human NPCs 35 days after transplantation stained with anti-human nuclei antibody (HuNu, green), NeuroTrace (fluorescent Nissl, blue) and counterstained with DAPI (grey). Scale bar: 50 um. Line graphs are plotted as mean ± sem. d: days, BLI: bioluminescence imaging, sbr = signal background ratio, ph: photons, sr: steradian

These data suggests that genetic immunosuppression is more reliable for graft survival, especially in long-term studies.

## Discussion

Immunodeficient mouse models are commonly used in preclinical stroke research to evaluate e.g., the efficacy of human cell-based therapies. As changes to the inflammatory responses after stroke may considerably alter overall stroke outcome, it is important to understand the stroke pathology of commonly used immunodeficient mouse models. Here, we identified distinct anatomical, gene expression and systemic changes in the stroke pathology between pharmacologically and genetically immunodeficient mice after stroke. These changes were most prominent in NSG mice that have deficiencies in the innate and adaptive immune system and less pronounced in Rag2^-/-^ and pharmacologically immunosuppressed mice with tacrolimus. The changes mainly affected the inflammatory and vascular responses after stroke. However, long-term graft survival was most consistent in NSG and Rag2^-/-^ mice.

A variety of cell types are responsible for postinjury inflammation in the brain after stroke. These cells may locally arise from resident microglia and astrocytes or are recruited from the peripheral blood circulation. Especially cells from the innate immune system such as neutrophils and monocytes have been extensively studied in experimental stroke [10]. It is assumed that these cells may predominantly be responsible for the secondary inflammatory injury by releasing proinflammatory factors, reactive oxygen species (ROS) and proteases [21]. Although neutrophils and monocytes constitute most of the remaining mouse immune cells in the immunodeficient NSG mice, dendritic cells and macrophages have been described to be detective because of the genetic NOD background [22]. We further observed an altered pattern in serum cytokines in NSG mice. These findings are directly in line with previous findings describing deficiencies of NSG mice in cytokines release [22]. These deficiencies in the cytokine and innate immune system may explain the strong deviation of the NSG gene expression profile after stroke to the other mouse models with intact innate immune systems. These differences also persisted after correcting for the genetic NOD background.

In contrast to the innate immune system, the role of the adaptive immune system after stroke is less clear. B and T lymphocytes are scarce in the CNS but can be found in the postischemic brain, which may be caused by cytokine signalling from the brain to the periphery or by the compromised blood-brain barrier after stroke. For instance, infiltration and activation of B and T lymphocytes in the brain have been associated with increased cognitive decline in an experimental stroke model and blocking adaptive immune responses has been suggested as a potential therapeutic target after stroke [23]. Both, the here used Rag2^-/-^ as well as tacrolimus immunosuppressed mice have deficiencies in the adaptive immune system and are lacking matured B- and T-lymphocytes (Rag2^-/-^) or respectively suppressed T-lymphocyte activation (tacrolimus). The lack of adaptive immune response caused a marked change in the brain tissue pathology as well as in the inflammatory and angiogenic gene expression signature. The main advantage in using pharmacological immunosuppression is that their time point of application can be chosen according to the experimental set-up. For instance, most studies using human cell-based strategies for stroke avoid the very acute phase for cell transplantations because of the hostile environment in the damaged brain regions; and choose transplantation around one week after stroke induction [3,24]. This set-up would facilitate to immunosuppress mice shortly before cell transplantation and allow a natural stroke progression for the first week. Tacrolimus has also been already successfully used in clinics for cell-based therapies in stroke and Parkinson’s and has therefore direct translational potential [25,26]. On the other hand, genetic mouse models are easier to handle and show generally a more consistent survival of the graft, as we have also observed in this study. Alternatively, future studies may use immune deficient mice such as NSG or Rag2^-/-^ Il2^-/-^ and CD47^-/-^ (Tripple KO) mice that are reconstituted with a humanised immune system and HLA-matched to the donor cells. Humanised mice may show a closer pathology to the WT [27], although these mice still have several limitations, including limited lifespan, incomplete human immune function and graft-versus-host disease etc [28].

Contrary to previous findings that described a reduced stroke volume of several immunodeficient mice [29,30], we did not observe anatomical changes in stroke outcome compared to immunocompetent WT mice. One reason for these inconsistencies may be the time point of stroke volume evaluation. Many studies evaluated stroke volume at 24 h after stroke induction to investigate acute effects of the stroke pathology. However, we were interested in long-term effects as regenerative cell therapies may have a therapeutic time course of weeks to months. At three weeks following stroke, major structural reorganisations occur, pushing the corpus callosum towards the brain surface, which in turn can affect the stroke volume estimates [31].

Findings from a growing number of preclinical and clinical studies suggest that post-stroke angiogenesis is a key restorative process for tissue preservation and repair, leading to improved outcome [14,15,32,33]. In our study, we observed marked upregulation of vascular repair in the peri-infarct regions of WT-Tacr, Rag2^-/-^ and NSG mice compared to immunocompetent WT mice 21 days after stroke. The improved vascular repair in histological brain sections was abolished if immunosuppressive tacrolimus was administered delayed at 7 dpi. As post-stroke angiogenesis starts from 3 dpi [34] and depends on the acute inflammatory and cytokine microenvironment in the injured brain, a delayed immunosuppression by 1 week seems to ensure that levels of vascular repair are more similar to that of control WT mice.

However, the pro-angiogenic responses in continuously immunosuppressed mice did not lead to improved functional outcome in our set-up. One explanation for the absence of improved functional recovery in immunodeficient mice could be that the observed angiogenesis only occurred transiently, as has been described previously [19,34]. The downregulation of angiogenesis-related genes at 21 dpi in the acutely tacrolimus immunosuppressed mice compared with mice receiving tacrolimus treatment 7 dpi further suggests the transient nature of post-stroke angiogenesis resulting from immunosuppression. Additionally, it is well known that the growth of new capillaries carries the risk for immature vessel formation, which may enhance haemorrhagic transformation and BBB damage; and mature vessels are associated with better functional outcomes [15,35]. We observed in all groups of animals a low pericyte coverage in the peri-infarct regions that tend to decrease especially in Rag2^-/-^animals, which had the highest vascular density in the injured brain regions. These changes in vascular responses following stroke are important to consider when evaluating the effects of cell-based therapies in immunosuppressed mice, as vascularization is often one of the important readouts to observe the effectiveness of a cell therapy [36,37].

One limitation of our study is that we used mice with different genetic backgrounds. Although we corrected the gene expression findings to the background of the mice, there might be confounding effects that are difficult to account for. For instance, mice with different backgrounds may have variable weight and size effects, time required to learn behavioral tasks or natural stroke recovery due to cerebral vascular architecture [38,39]. We used two cohorts of animals for the gene expression and anatomical studies after stroke, which may add to the variability of the data. Future studies may profit from recent advances in spatially resolved transcriptomics approaches that could directly associate gene expression signatures to anatomical changes in the same animal [40]. However, the natural temporal delay in gene expression and anatomical and functional recovery may complicate understanding of complex processes in vivo, such as whether angiogenesis occurs transiently after stroke. The pharmacological immunosuppression with tacrolimus occurred via osmotic mini pumps that ensured a constant delivery rate previously validated in other studies [41–43]; however, we do not know the spatiotemporal distribution and delivery efficacy of tacrolimus to the injured brain regions, an irregular delivery may influence the survival of the grafted cells. Furthermore, tacrolimus may additionally influence the survival of the graft independent of the calcineurin-pathway [44]. Future studies may compare different routes of delivery for tacrolimus and alternative immunosuppressive regimes to optimize graft survival.

In sum, our study identifies key changes in anatomy, physiology, and gene expression of popular immunosuppressed mouse models after stroke. These changes in stroke progression should be considered when investigating molecular mechanisms of stroke or evaluating therapies that require immunosuppression.

## Materials and Methods

### Experimental design

Stroke pathology was compared between (1) wildtype (WT) C57BL/J mice with three immunosuppressant mouse models: (2) WT C57BL/J mice that were immunosuppressed with tacrolimus (3) Rag2^-/-^ mice and (4) NSG mice. All mice received a large photothrombotic stroke to the right sensorimotor cortex and underwent regular behavioral tests at baseline, 3,7,14,21 days after stroke induction. At 21 days after stroke, mice from each group were divided into two subgroups: Brains from the first subgroup were collected, fixed and histologically analysed; brains from the other group were microdissected around the stroke region and prepared for RNAseq analysis. To test the effects of delayed tacrolimus treatment, we compared between (1) wildtype C57BL/J mice with (2) WT C57BL/J mice that were acutely immunosuppressed mice with tacrolimus (0 dpi) and (3) WT C57BL/J mice that received tacrolimus delayed at 7 dpi. The brains were treated equivalently to the previous set-up to obtain comparable histology and RNAseq data.

Long-term survival and successful immunosuppression were tested by local intraparenchymal transplantation of human NPCs to 1) WT C57BL/J mice (2) WT C57BL/J mice that were immunosuppressed with tacrolimus (3) Rag2^-/-^ mice and (4) NSG mice. Luciferase expressing NPCs were tracked over 35 days using *in vivo* bioluminescence imaging and brain tissue was collected to identify the transplanted cells in brain sections.

### Animals

All procedures were conducted in accordance with governmental, institutional (University of Zurich), and ARRIVE guidelines and had been approved by the Veterinarian Office of the Canton of Zurich (license: 209/2019). In total, 20 WT (WT) mice (histology: 7, RNAseq: 8, cell therapy: 5) with C57BL/6 background mice, 17 tacrolimus treated WT mice (histology:8, RNAseq: 8, cell therapy: 5), 19 Rag2^-/-^ mice (histology: 5, RNAseq: 9, cell therapy: 5) and 19 NSG mice (histology: 10, RNAseq: 4, cell therapy: 5) were used. Mice that were used for histology and RNAseq both performed the behavioral tasks. We used both sex female and male and mice were 3 months of age. Mice were housed in standard type II/III cages on a 12h day/light cycle (6:00 A.M. lights on) with food and water ad libitum. All mice were acclimatized for at least a week to environmental conditions before set into experiment.

### Photothrombotic lesion

Anaesthesia was performed using isoflurane (5% induction, 1.5-2% maintenance, Attane, Provet AG) and adequate sedation was confirmed by tail pinch. Novalgin (1mg/ml) was applied via drinking water; 24 h prior to the procedure and for three consecutive days directly after stroke surgery. Cerebral ischemia was induced by photothrombotic stroke surgery as previously described (Labat-gest and Tomasi, 2013; Rust et al., 2019b). Briefly, animals were fixed in a stereotactic frame (David Kopf Instruments), the surgical area was sanitized, and the skull was exposed through a cut along the midline.

A cold light source (Olympus KL 1,500LCS, 150W, 3,000K) was positioned over the right forebrain cortex (anterior/posterior: −1.5–+1.5 mm and medial/lateral 0 mm to +2 mm relative to Bregma). Rose Bengal (15 mg/ml, in 0.9% NaCl, Sigma) was injected intraperitoneally 5 min prior to illumination and the region of interest was subsequently illuminated through the intact skull for 12 min. To restrict the illuminated area, an opaque template with an opening of 3 × 4 mm was placed directly on the skull. The wound was closed using a 6/0 silk suture and animals were allowed to recover.

### Blood Perfusion by Laser Doppler Imaging

Cortical perfusion was evaluated using Laser Doppler Imaging (Moor Instruments, MOORLDI2-IR). Briefly, animals were fixed in a stereotactic frame and the region of interest was shaved and sanitized. To expose the skull, a cut was made along the midline and the brain was scanned using the repeat image measurement mode. All data were exported and quantified in terms of total flux in the ROI using Fiji (ImageJ).

### Tacrolimus pump implantation

For pharmacological immunosuppression, Alzet osmotic pumps (Model 1004, Cupertino, CA, USA) filled with 100μl tacrolimus solved in a mixture of DMSO and PEG 300 (50mg/ml; 7.56mg/ml concentration) were implanted subcutaneously on the back according to the manufacturer’s protocol. Briefly, animals were anesthetized using isoflurane (5% induction, 1.5-2% maintenance, Attane, Provet AG), the surgical area was sanitized and shaved. The implantation site was exposed through a midscapular incision. An implantation pocket was created using a hemostat. The pump was inserted into the pocket and the wound was closed with sutures.

### Tissue processing and immunofluorescence

Animals were euthanized using pentobarbital (i.p, 150 mg/kg body weight, Streuli Pharma AG) and perfused transcardially with Ringer solution (containing 5 ml/l Heparin, B. Braun) followed by paraformaldehyde (PFA, 4%, in 0.2 M phosphate buffer, pH 7). Brain tissue was collected and postfixed for 6 h in 4% PFA. For cryoprotection, tissue was transferred to 30% sucrose and stored at 4°. Coronal sections were cut at a thickness of 40 μm using a sliding microtome (Microm HM430, Leica), collected, and stored as free-floating sections in cryoprotectant solution at −20°C.

For immunostaining, brain sections were blocked with 5% normal donkey serum for 1 h at room temperature and incubated with primary antibodies (rabbit anti-GFAP 1:200, Dako; goat anti-Iba1, 1:500 Wako; NeuroTrace™ 1:200, Thermo Fischer; mouse anti-NeuN Antibody 1:500, Merck, #MAB377; rabbit anti-Neurofilament 200 antibody 1:200, Merck, #N4142; guinea pig anti-Neurofilament L, 1:200, Synaptic Systems, rat anti-CD31 antibody 1:50, BD Biosceicnes; goat anti-CD13, 1:200; R&D Systems) overnight at 4°C. The next day, sections were incubated with corresponding secondary antibodies (1:500, Thermo Fischer Scientific). Nuclei were counterstained with DAPI (1:2,000 in 0.1 M PB, Sigma). Mounting was performed using Mowiol.

### Vascular quantification

Vascular parameters (vessel area fraction, length, branching and nearest distance) were identified using an automated ImageJ script as previously described for brain and retinal vasculature [45,46]. Briefly, for total vessel area fraction, the area covered by anti-CD31 staining was quantified using ImageJ after applying a constant threshold. The vascular length was evaluated using the“skeleton length” tool and number of branches was calculated with the “analyse skeleton” tool. Results were normalized per mm^2^ of brain tissue for vascular length and number of branches.

### EdU administration

5-ethynyl-2’-deoxyuridine (EdU, 50 mg/kg body weight, ThermoFischer) was applied intraperitoneally on day 7 after stroke to label proliferating vascular endothelial cells. EdU incorporation was detected 21 days after stroke using the Click-it EdU Alexa Fluor 647 Imaging Kit (ThermoFischer) on 40 μm coronal sections.

### Blood plasma multiplex cytokine analysis

Blood was collected through the tail-vein at baseline, 7- and 21-days post injury. Blood plasma cytokines levels (of IFN-γ, IL-1β, IL-2, IL-4, IL-5, IL-6, KC/GRO, IL-10, IL-12p70, and TNF-α) were measured using the Proinflammatory Panel 1 Mouse kit, a chemiluminescence-based assays from Meso Scale Discovery (MSD, Gaithersburg, MD, USA) according to the manufacturer’s protocol. Analyses were done using a QuickPlex SQ 120 instrument (MSD) and DISCOVERY WORKBENCH^®^ 4.0 software.

### Gene expression analysis

Total RNA of stroked cortical tissue and the corresponding contralesional site was isolated using TRIzol extraction followed by the RNeasy RNA isolation kit (Qiagen) including a DNase treatment to digest residual genomic DNA. All samples had a RIN value of >8.5.

Library preparation, sequencing, read processing, read alignment, and read counting was performed at the Functional Genomics Center Zurich (FGCZ) core facility. For library preparation, the TruSeq stranded RNA kit (Illumina Inc.) was used according to the manufacturer’s protocol. The mRNA was purified by polyA selection, chemically fragmented and transcribed into cDNA before adapter ligation. Analysis of RNA sequencing data and gene set enrichment analysis was performed using EdgeR [47] and clusterProfiler 4.0 [48].

Transcriptomic raw data are available to readers via Gene Expression Omnibus with identifier GSE208605.

### Behavioral studies

Animals were tested on the (1) runway, (2) ladder rung test and (3) the rotarod test. All animals were tested 3 days before surgery to establish baseline performance and 3, 7, 14 and 21 days after stroke induction. Performance on the runway and the ladder rung was evaluated using a recently established deep learning-based protocol [16].

Briefly, a runway walk was performed to assess whole body coordination during overground locomotion. Mice were recorded crossing the runway with a high-definition video camera (GoPro Hero 7) at a resolution of 4000 × 3000 and a rate of 60 frames per second from three perspectives. Each animal was placed individually on one end of the transparent plexiglas basin and was allowed to walk for 3 minutes.

The ladder rung test was performed using the same set-up as for the runway to assess skilled locomotion; with the only difference that the runway was replaced with a horizontal ladder on which the spacing of the rungs is variable. The animals were trained to cross the ladder without external reinforcement. A total of at least three runs per animal and per session were recorded. A step was defined as misstep when the toe tips of the animal reached a threshold of >0.5 cm below the ladder height.

The rotarod test is a standard sensory-motor test to investigate the animals’ ability to stay and run on an accelerated rod (Ugo Basile, Gemonio, Italy). All animals were pre-trained to stay on the accelerating rotarod (slowly increasing from 5 to 50 rpm in 300s) until they could remain on the rod for > 60 s. During the performance, the time and speed was measured until the animals fell or started to rotate with the rod without running. The test was always performed three times and means were used for statistical analysis. The recovery phase between the trials was at least 10 min.

### Cell transplantations

Induced pluripotent stem cell (iPSC)-derived neural progenitor cells (NPCs) were generated as previously described [20]. In brief, NPCs at passage number < 11 were used in all experiments. Intraparenchymal cell transplantation was performed as previously described [49]. In brief GFP^+^/Luc^+^ NPCs were thawed, washed, and diluted to a final concentration of 8 × 10^4^ cells/μL in sterile 1x PBS (pH 7.4, without calcium or magnesium; Thermo Fisher Scientific). Cells were stored on ice until transplantation. Mice were anesthetized using isoflurane (5% induction, 1.5% maintenance: Attane, Provet AG). Analgesic (Rimadyl; Pfizer) was administered subcutaneously prior to surgery (5 mg/kg body weight). Animals were fixed in a stereotactic frame (David Kopf Instruments), the surgical area was sanitized, and the skull was exposed through a midline skin incision to reveal lambda and bregma points. Coordinates were calculated (the coordinates of interest chosen for this protocol were: AP: + 0.5, ML: + 1.5, DV: −0.8 relative to bregma) and a hole was drilled using a surgical dental drill (Foredom, Bethel CT). Mice were injected with 1.6 × 10^5^ cells (2μL/injection) using a 10-μL Hamilton microsyringe (33-gauge) and a micropump system with a flow rate of 0.3 μL/min (injection) and 1.5 μl/min (withdrawal). The wound was closed with suture.

### *In vivo* bioluminescence imaging and analysis

Bioluminescence imaging (BLI) experiments were performed with the IVIS Spectrum CT (PerkinElmer). Animals were imaged in regular intervals starting 2h after transplantation for up to 35 days. 30mg/ml D-luciferin potassium salt (PerkinElmer) was dissolved in 1x PBS (pH 7.4, without calcium or magnesium; Thermo Fisher Scientific) and sterilized through a 0.22 μm syringe filter. Luciferin was injected intraperitoneally with a final dose of 300 mg/kg body weight before isoflurane anaesthesia (5% induction, 1.5% maintenance; Attane, Provet AG). The head region was shaved before imaging. Image acquisition was performed for 20 min using the following settings: Field of View: A, Subject height: 1.5 cm, Binning: 16, F/Stop: 2. Exposure time (ranging from 1s to 60s) was set automatically by the system to reach the most sensitive setting. Imaging parameters and measurement procedures were kept consistent between mice.

The regions of interest (ROIs) were manually drawn based on anatomical landmarks (eyes, ears and snout) on each image. Plotting and statistical analysis was performed using RStudio. The brain-specific signal was calculated and corrected for nonspecific signal taken from a ROI on the skin of the animal’s back and for noise taken from a ROI outside the mouse. Signal-to-noise ratio (SNR) was calculated by dividing the total photon flux (ph/s/cm^2^/sr) by the standard deviation of the noise. Bioluminescent signal was calculated by subtracting the background flux from the total photon flux per ROI.

### Statistical Analysis

Statistical analysis was performed using RStudio (4.04 (2021-02-15). Sample sizes were designed with adequate power according to our previous studies [14,50,51] and to the literature [3,15]. All data were tested for normal distribution by using the Shapiro-Wilk test. Multiple comparisons (NSG, Rag2^-/-^, WT-Tacr, WT) were initially tested for normal distribution with the Shapiro-Wilk test. The significance of mean differences between normally distributed multiple comparisons was assessed using repeated measures ANOVA with post-hoc analysis (p adjustment method = holm). Normally distributed data (WT-Tacr_acute_ vs. WT-Tacr_delayed_) were tested for differences with a two-tailed unpaired one-sample *t-*test to compare changes between the two groups. Variables exhibiting a skewed distribution were transformed, using natural logarithms before the tests to satisfy the prerequisite assumptions of normality. Data are expressed as means ± SD, and statistical significance was defined as \**p* < 0.05, \*\**p* < 0.01, and \*\*\**p* < 0.001.

## Data availability

Transcriptomic raw data are available to readers via Gene Expression Omnibus with identifier GSE208605. All other raw data are available upon request.

## Acknowledgements

The authors thank the Functional Genomics Center Zurich (FGCZ) for support with the RNASeq data.

