## Supplementary material for "Molecular and anatomical roadmap of stroke pathology in immunodeficient mice": Suppl. Material

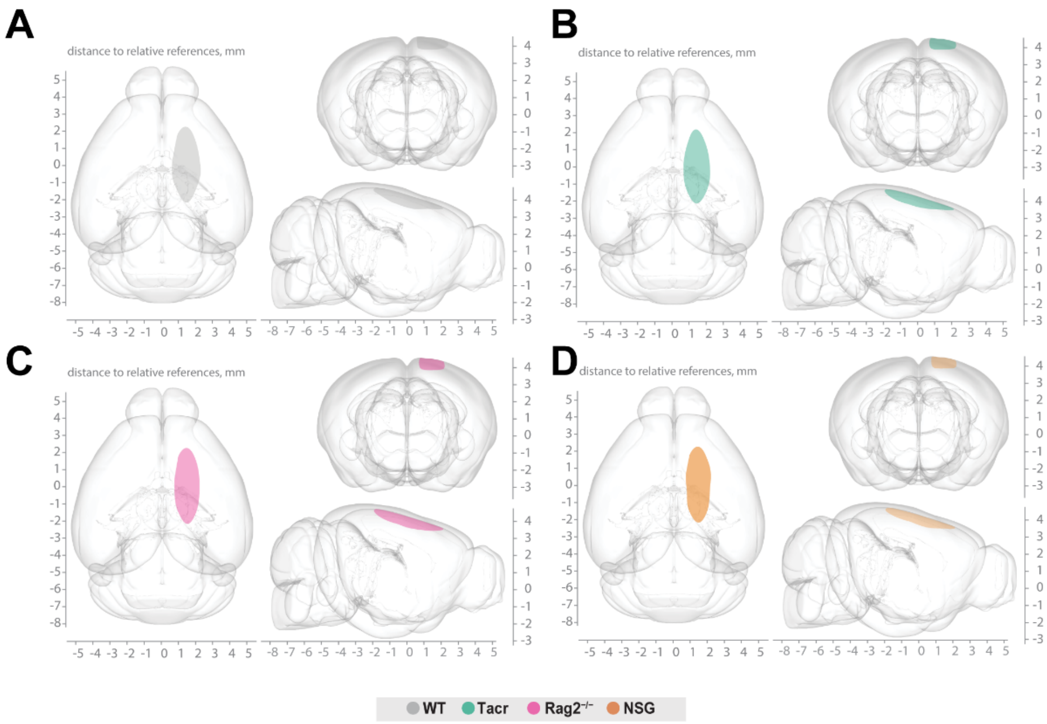


**Suppl. Fig. 1: Stroke volume and location**. 3D reconstruction of stroke location within a brain template from three perspectives of (A) C57BL/6J wildtype (WT), (B) Tacrolimus immunosuppressed wildtype (WT-Tacr), (C) recombination activating gene 2 deficient mice ($Rag2^{-/-}$) and (D) NOD scid gamma mice (NSG).

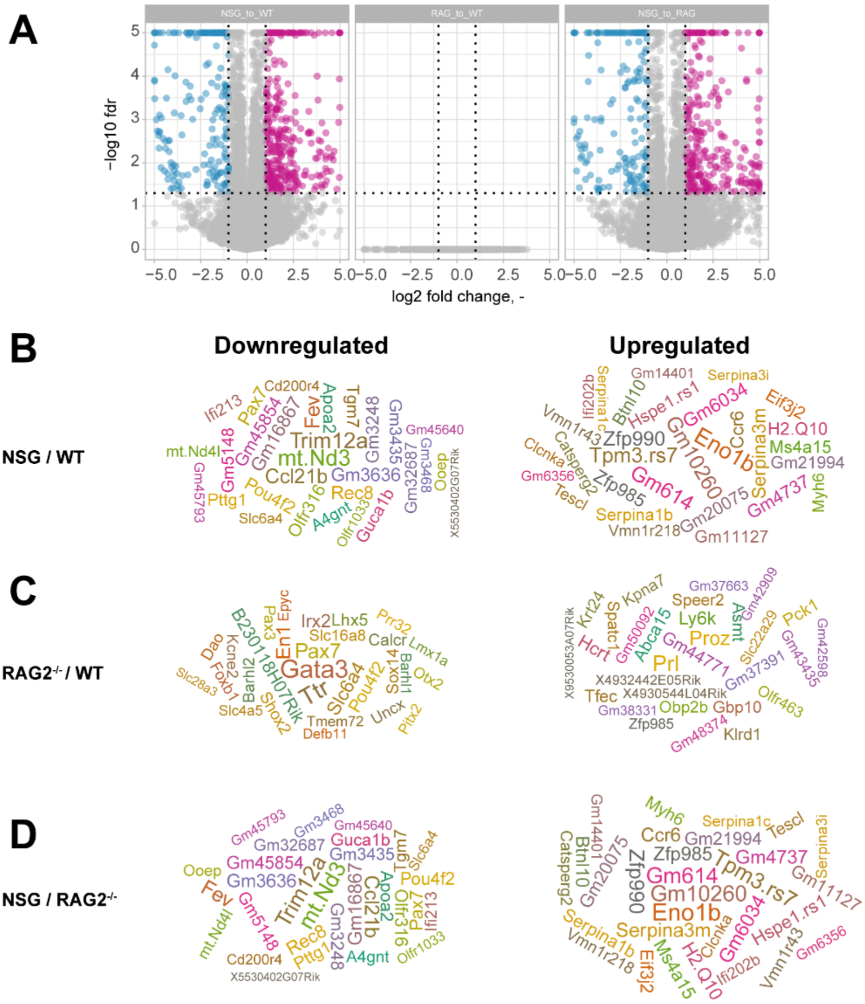


**Suppl. Fig. 2: Gene expression differences of non-stroked WT and non-stroked immunosuppressed mice.** (A) Volcano plot of non-stroked NSG mice to non-stroked WT (left) non-stroked Rag2$^{-/-}$, to non-stroked WT (middle) and non-stroked NSG to non-stroked Rag2$^{-/-}$ (right) (B) List of top 30 upregulated and downregulated genes after stroke in (B) non-stroked NSG mice to non-stroked WT (C) non-stroked Rag2$^{-/-}$, to non-stroked WT and (D) non-stroked NSG to non-stroked Rag2$^{-/-}$.

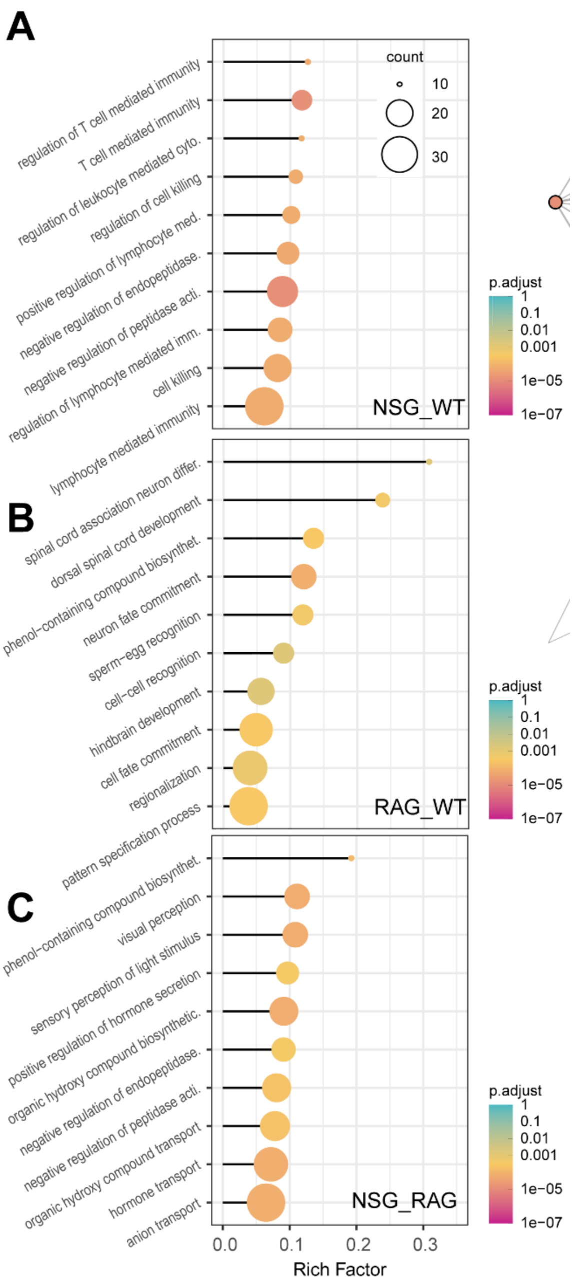


**Suppl. Fig. 3: Gene set enrichment analysis in stroked mice.** Lollipop plot and of top 10 pathways enriched in (A) non-stroked NSG mice to non-stroked WT (B) non-stroked Rag2$^{-/-}$, to non-stroked WT and (C) non-stroked NSG to non-stroked Rag2$^{-/-}$. Size of dots represents the number of genes in the pathway and color of dots represents adjusted p value.

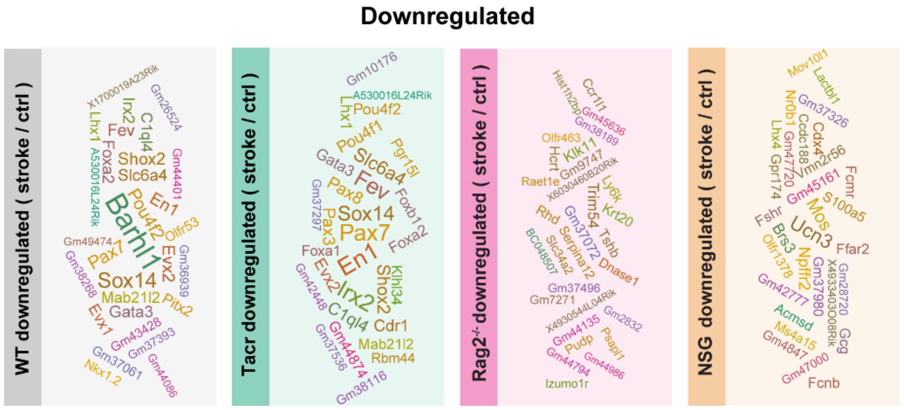


**Suppl. Fig. 4: List of top 30 downregulated genes after stroke in the respective group of immunodeficient mice.**

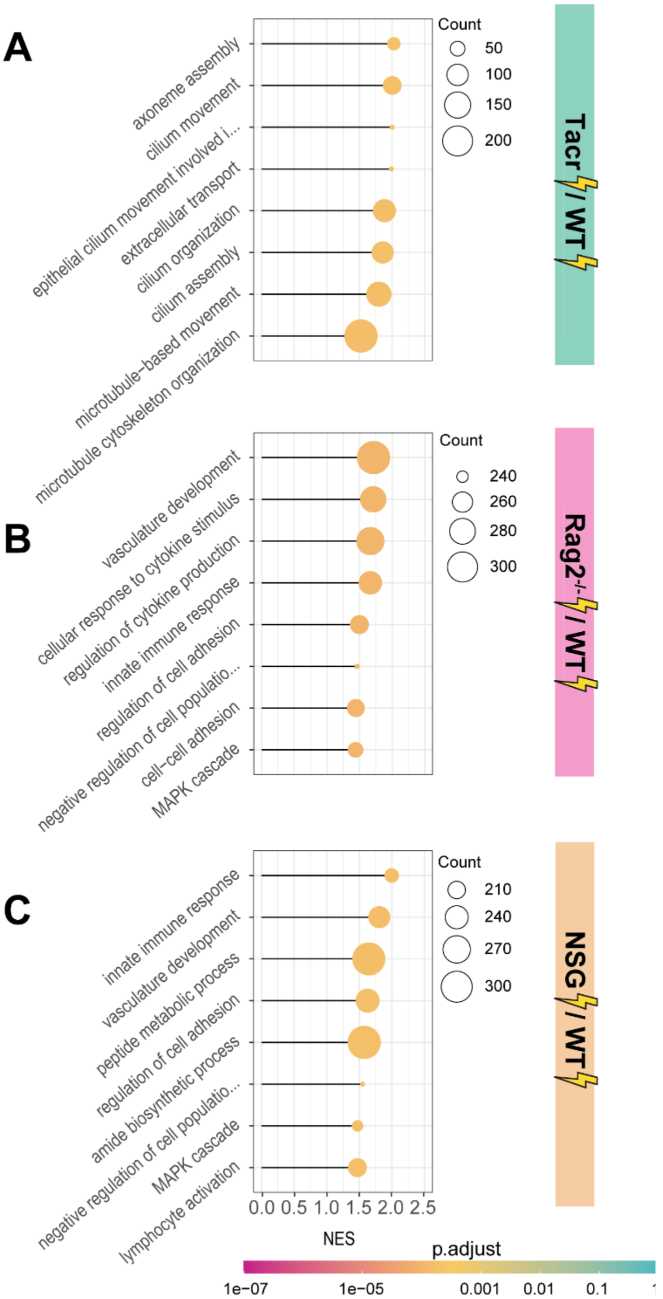


**Suppl. Fig. 5: Gene set enrichment analysis in stroked mice.** Lollipop plot and of top 10 pathways enriched in (A) stroked WT-Tacr mice to stroked WT (B) stroked Rag2$^{-/-}$, to stroked WT and (C) stroked NSG to stroked WT$^{-/-}$. Size of dots represents the number of genes in the pathway and color of dots represents adjusted p value.

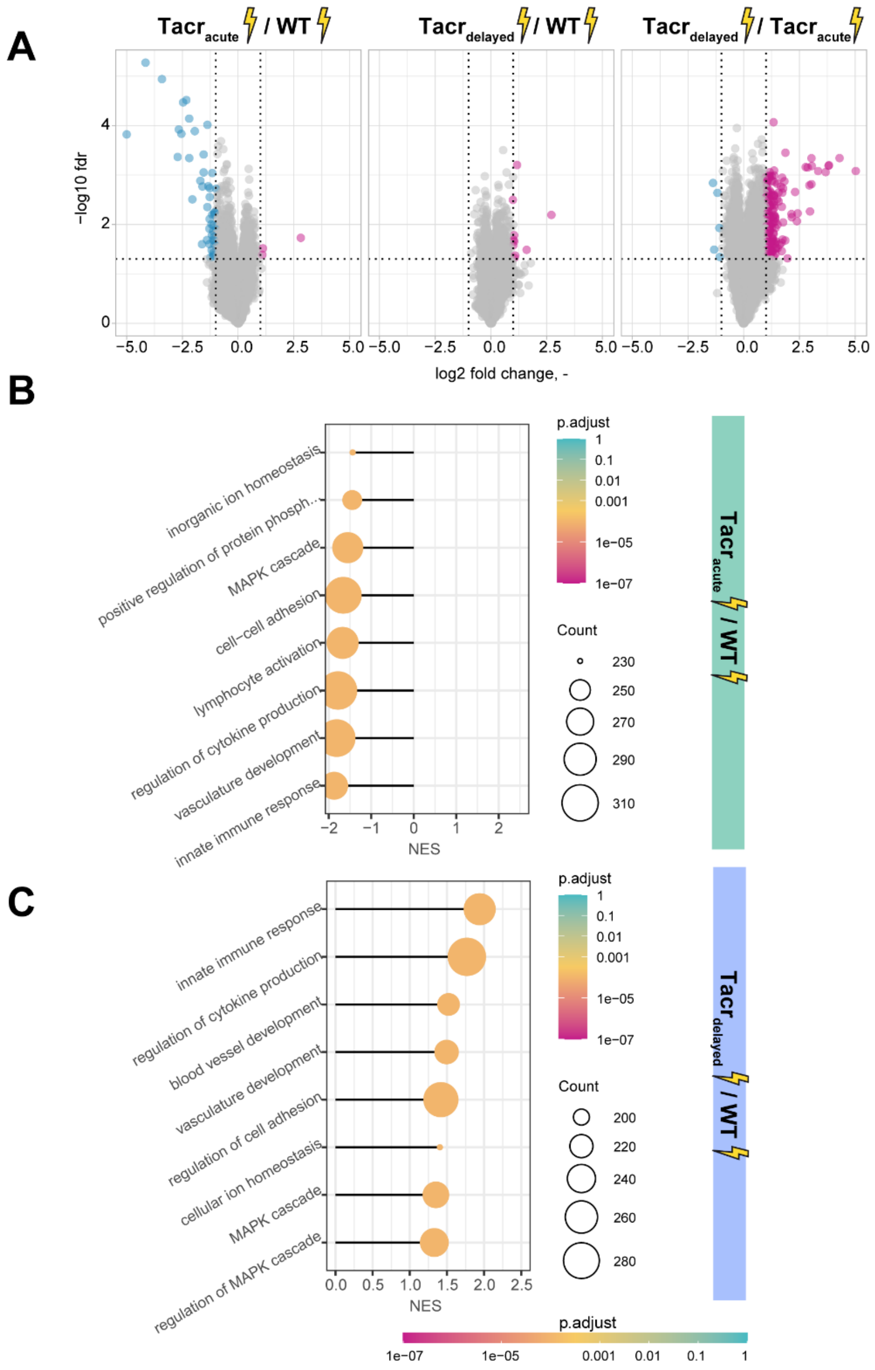


**Suppl. Fig. 6: Gene expression differences between acute and delayed Tacrolimus treatment after stroke. (A)** ) Volcano plot of stroked WT-Tacr$_{acute}$ mice to stroked WT (left) stroked WT-Tacr$_{delayed}$ mice to stroked WT (middle) and stroked WT-Tacr$_{delayed}$ mice to stroked WT-Tacr$_{acute}$ (right). **(B)** Lollipop plot and of top 10 pathways enriched in (A) stroked WT-Tacr$_{acute}$ mice to stroked WT and (B) stroked WT-Tacr$_{delayed}$ mice to stroked WT. Size of dots represents the number of genes in the pathway and color of dots represents adjusted p value.
